# A potent neutralizing mouse monoclonal antibody specific to dengue virus type 1 Mochizuki strain recognized a novel epitope around the N-67 glycan on the E protein: a possible explanation of dengue virus evolution regarding the acquisition of N-67 glycan

**DOI:** 10.1101/2020.10.03.324780

**Authors:** Tomohiro Kotaki, Atsushi Yamanaka, Eiji Konishi, Masanori Kameoka

## Abstract

Analysis of the neutralizing epitope of dengue virus (DENV) is important for the development of an effective dengue vaccine. A potent neutralizing mouse monoclonal antibody named 7F4 was previously reported and, here, we further analyze the detailed epitope of this antibody. 7F4 recognized a novel conformational epitope close to the N-67 glycan on the E protein. This antibody was specific to the DENV that lacks N-67 glycan, including the Mochizuki strain. Interestingly, the Mochizuki strain acquired N-67 glycan by 7F4 selective pressure. DENVs might have evolved to escape from this antibody considering that most of currently circulating DENVs possess N-67 glycan. However, this suggests that 7F4 epitope might be useless as a vaccine target. Nevertheless, this study demonstrated the existence of epitopes competing for 7F4 epitope, which are involved in neutralization. This study describes the importance of antibodies recognizing epitopes near the N-67 glycan for future dengue vaccine development.

## 1. Introduction

The genus *Flavivirus* of the family *Flaviviridae* includes four types of dengue virus (dengue type 1 to 4 viruses; DENV-1 to DENV-4) (Pierson and Diamond, 2013). DENV causes dengue and severe dengue, which are globally important arboviral diseases (WHO, 2012). An estimated 390 million cases of dengue infection occur annually worldwide (Bhatt et al., 2013), out of which 100 million develop symptomatic dengue and 21,000 die (Thomas and Endy, 2013). Dengue vaccine (CYD-TDV) was approved for use in several countries (Hadinegoro et al., 2015; Aguiar et al., 2016); however, its efficacy is still low especially for sero-negative individuals (Aguiar et al., 2016). Further analysis of antibodies induced by DENV infection and their neutralization mechanism may help develop a more effective vaccine against dengue.

DENV contains a single-stranded, positive-sense RNA. The genomic viral RNA is translated in the cytoplasm as a single polyprotein, which is further processed by the host and viral proteases into three structural proteins and seven nonstructural proteins. The capsid (C), precursor of the membrane (prM), and envelope (E) comprise the structural proteins. DENV assembly takes place in the endoplasmic reticulum (ER) (Lorenz et al., 2003, 2004). The C-RNA nucleocapsid complex buds on the surface of the ER lumen and then acquires a lipid bilayer containing 180 copies of prM-E heterodimers.

At the mildly low pH of the trans-Golgi network, the prM-E heterodimers are rearranged to 90 E-dimers, followed by cleavage of the pr protein from prM on the viral particle (Yu et al., 2008). Thus, the viral particle acquires infectivity (viral maturation). Virus particles exhibit conformational dynamics referred to as viral “breathing” (Kuhn et al., 2015). The released infectious virus enters the cell by clathrin-mediated endocytosis and is then trafficked to the late endosomes. The acidic pH conditions in the late endosome rearrange the virion structure from E-dimer to E-trimer, resulting in the exposure of fusion loop regions, and membrane fusion (Modis et al., 2003).

The E protein is a major surface protein which contains most of the neutralizing epitopes, and thus, is a major target as a vaccine antigen. The E protein structure consists of three domains - I, II, and III (Modis et al., 2003). Domain I (DI) is an N-terminal domain structurally located at the ectodomain center. Domain II (DII) is a finger-like structure domain which is involved in stabilizing the E protein dimer and contains the fusion loop region (Allison et al., 2001). Domain III (DIII) is an Ig-like domain containing the putative receptor binding motif (Crill and Roehrig, 2001; Rey et al., 1995).

The E protein of most flaviviruses has one or two N-linked glycosylation motifs (N-X-T/S, X could be any amino acid except P) at N-153/154 and/or N-67. Loss of glycan is not critical for viral growth since some naturally circulating flaviviruses lack these N-linked glycans (Vorndam et al., 1993; Roehrig et al., 1998; Scherret et al., 2001). The glycosylation at N-153/154 located on DI plays a vital role in both infectivity and assembly and, thus, is highly conserved among the flaviviruses (Zhang et al., 2017; Mossenta, 2017). On the other hand, glycosylation at N-67 located on DII is unique to DENV. This glycan is associated with virus attachment to the C-type lectin DC-SIGN, expressed on dendritic cells and monocyte/macrophages (Modis et al., 2005; Mondote et al., 2007; Liang et al., 2018), determining tropism and pathogenesis of DENV (Wu et al., 2000). Furthermore, the glycan is associated with virion release (Lee et al., 2010). Interestingly, DENV-1 Mochizuki strain, which is the oldest DENV, isolated in 1943, lacks glycosylation at this site (Hotta, 1952; Ishak et al., 2001).

Antibody epitope is associated with neutralizing potency. Generally, weak neutralizing antibodies target the fusion loop epitope on DII (Dejnirattisai et al., 2015). These antibodies are cross-reactive to all four serotypes of DENV and inhibit only viral attachment. Meanwhile, potent neutralizing antibodies target E-dimer or quaternary-structure epitopes, the hinge region of DI-DII, or DIII. (Teoh et al., 2012; de Alwis et al., 2012; Messer et al., 2014; Dejnirattisai et al., 2015, Wong et al., 2017, Hu et al., 2019). These antibodies tend to exhibit limited cross-reactivity, and inhibit both pre- and postviral attachment.

We previously reported a mouse monoclonal antibody (D1-IV-7F4, or 7F4) displaying a 100 to 1,000-fold stronger neutralizing activity than other anti-dengue mouse antibodies (Yamanaka et al., 2013). The deduced epitope of 7F4 includes the 118th amino acid on DII, based on an escape mutant generated in the study, and it was distinct from the other known neutralizing epitope (Yamanaka et al., 2013). This antibody is worth analyzing in detail in order to develop an effective vaccine against dengue. In this study, we conducted a detailed epitope mapping of 7F4, by generating neutralization-escape mutants and by applying the site-directed mutagenesis method. The epitope was identified to the region near the N-67 glycan. This study demonstrated the involvement of 7F4 antibody selection pressure on viral evolution and the importance of this epitope in the development of a dengue vaccine.

## 2. Results

### General characteristics of 7F4

The main findings of the previous study were that 1) 7F4 was specific to DENV-1, 2) NT_50_ to DENV-1 was approximately 0.02 μg/mL, and 3) the epitope included the 118^th^ amino acid on E protein (Yamanaka et al., 2013). We further analyzed the neutralization mechanism, specificity, and the epitope of 7F4.

A time of addition assay demonstrated that 7F4 inhibited both pre- and postviral attachment, indicating that 7F4 blocked both viral attachment and viral conformational change (Fig. 1). 7F4 neutralized DENV-1 Mochizuki strain, but not the other DENV-1, including the Hawaii strain and Indonesian clinical isolate, indicating narrow specificity (Fig. 1) (Sjatha et al., 2012). 7F4 was reactive to detergent-inactivated E protein by ELISA, but not reactive to reduced and heat-inactivated E protein by Western blot (Table 1). These results suggest that 7F4 recognized the conformational E-monomer epitope in the Mochizuki strain. Table 1 shows a summary of the characteristics of 7F4 compared to 4G2 (weak neutralizing mouse monoclonal antibody targeting fusion loop) (Fig. S1).

**Figure 1.**
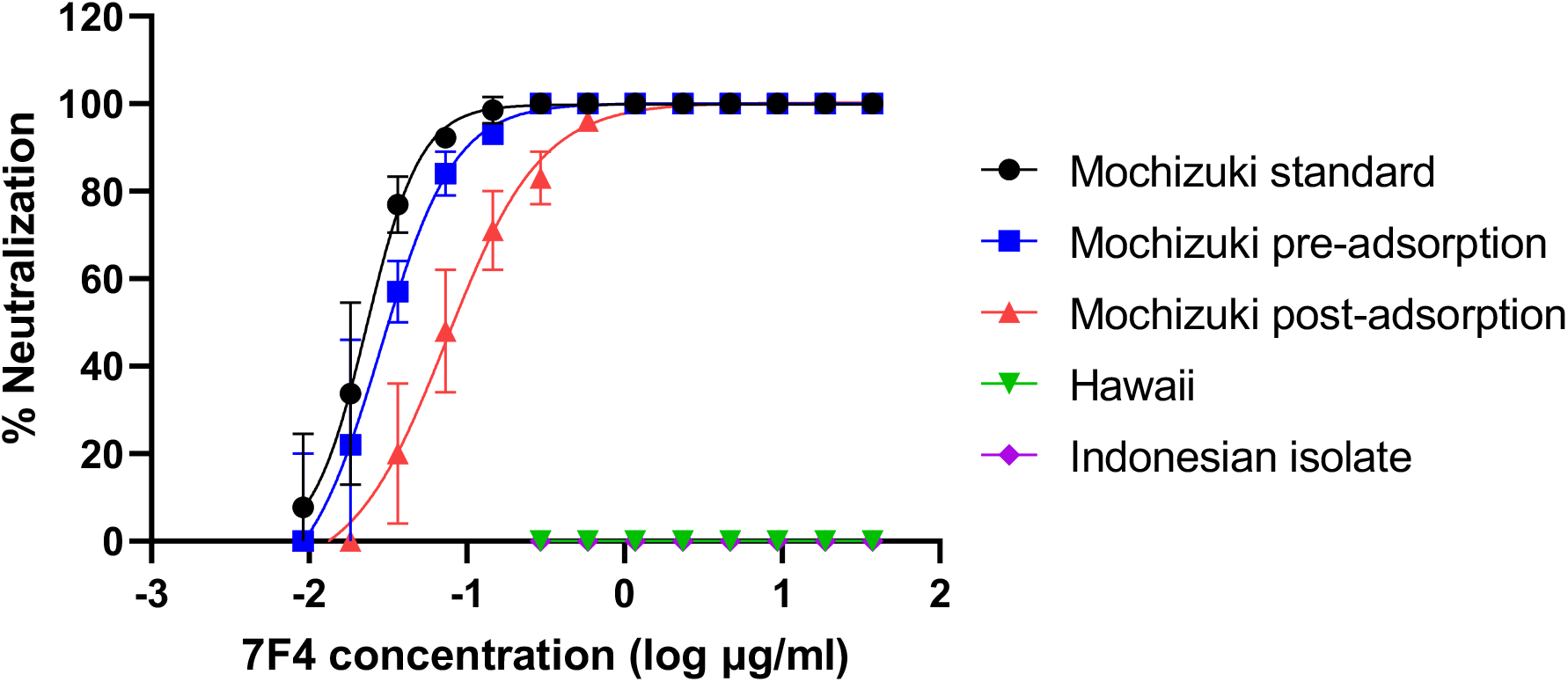
Neutralizing activity of 7F4. Results of the time of addition assay and neutralization test against Hawaii and Indonesian isolate are shown. The averages and SDs of the two independent experiments are shown.

**Table 1.** General characteristics of 7F4.

|  | Cross-reactivity | NT<sub>50</sub> (µg/mL) |  |  | Used as a detector antibody |  |
| --- | --- | --- | --- | --- | --- | --- |
|  |  | Standard method | Preattachment | Postattachment | ELISA* | Western blot ** |
| 7F4 | Mochizuki specific | 0.02 | 0.03 | 0.07 | Reactive | Nonreactive |
| 4G2 | Pan-flavivirus | 16.7 | 57.1 | >100 | Reactive | Reactive |

\*The virus was inactivated by the detergent.
\*\*The virus was heat-inactivated in reduced conditions.

### Generation of the neutralization-escape mutants of Mochizuki strain to 7F4

Neutralization-escape mutants of Mochizuki strain to 7F4 were further generated with various antibody concentrations in Vero or C6/36 cells in order to identify the detailed 7F4 epitope. The prM-E sequences were compared with the wild type (WT) Mochizuki strain, escape mutants, and passage control virus. There was no difference in the prM-E sequences between the passage control and WT Mochizuki strain. Escape mutants obtained in the Vero cells acquired several kinds of mutations, which were divided into three groups (Table 2). Group 1 possessed a mutation at the 1,286^th^ nucleotide of the Mochizuki strain (Genbank accession No. AB074760; A to G) translated to K118E in the E, as reported previously (Yamanaka et al., 2013). Group 2 had a mutation at the 1,133^th^ nucleotide (A to G), translated to N67D in the E. Group 3 had simultaneous mutations at the 1,106^th^ (G to A) and 1,140^th^ nucleotides (T to C), translated to E58K and I69T in the E, respectively. Furthermore, a nucleotide mutation at the 475^th^ nucleotide (G to A) translated to M13I in the prM was introduced in all 3 groups. All escape mutants were confirmed to be resistant to 7F4; the escape mutants were 1450-fold resistant compared to those of the WT Mochizuki strain (Table 2). In contrast to the Vero cells, all the escape mutants derived from C6/36 cell had only two mutations: M13I in prM and I69T in E (designated as group 4) (Table 2). Interestingly, E58K mutation was not observed in the escape mutant derived from C6/36 cells.

These mutations were plotted on the ribbon diagram of DENV-1 prM-E proteins (Fig. 2). The mutations in E were located in the central portion of DII. Interestingly, mutation I69T completed the N-linked glycosylation motif in the Mochizuki strain (N_67_-I_68_-I_69_ to N_67_-I_68_-T_69_) resulting in glycosylation at N-67 (Table 2). The Mochizuki strain, which originally possessed only one glycan at N-153, acquired a second glycan at N-67 resulting from neutralization-escape to 7F4. Glycosylation of the escape mutants was confirmed by Western blot (Fig. S2).

**Figure 2.**
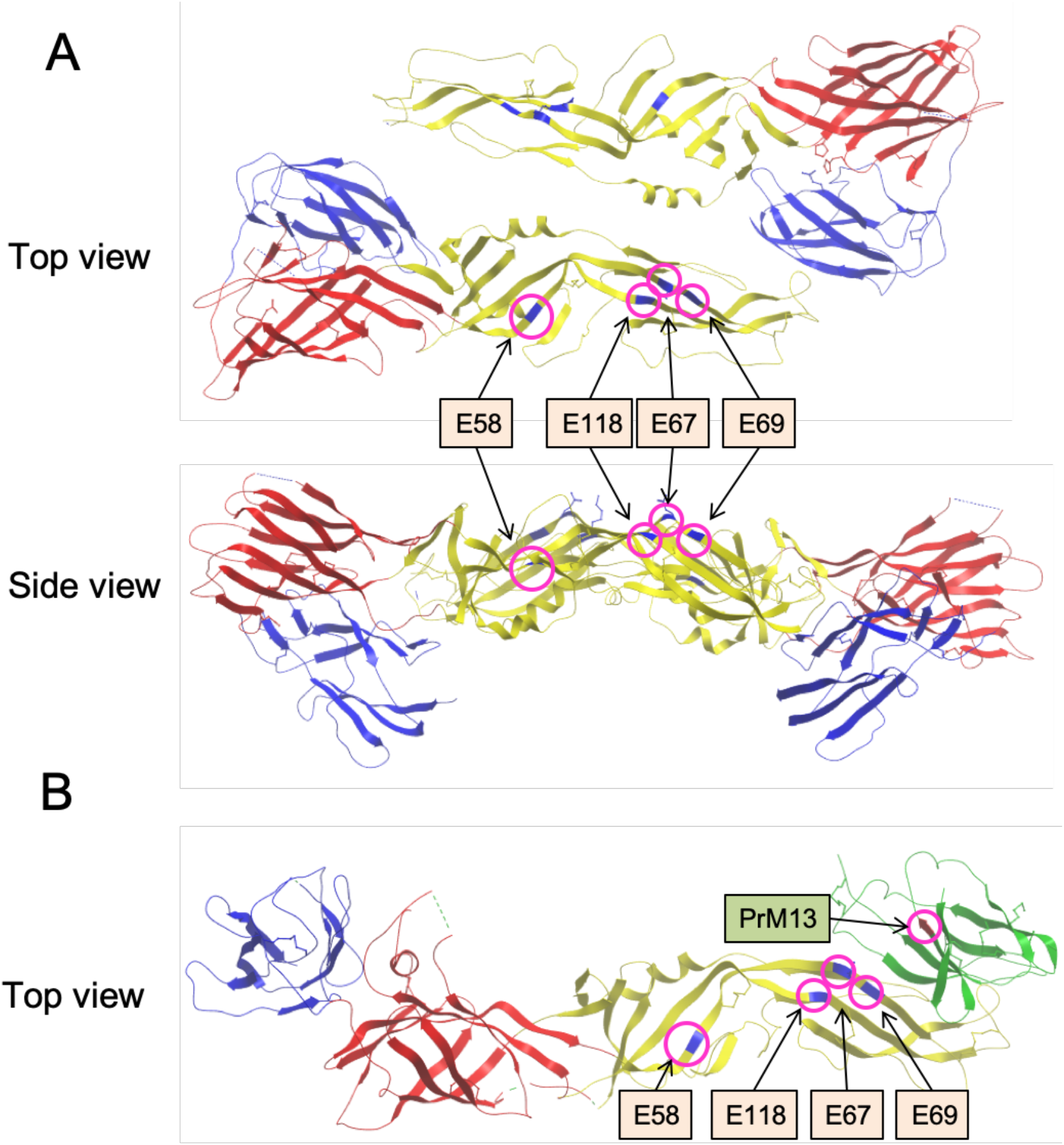
Amino acid mutations introduced in the escape mutants to 7F4. (A) Positions of the amino acid mutations on a DENV E protein dimer. The ribbon diagram is based on Protein Data Bank (PDB) accession No. 4GSX. EDI, EDII, and EDIII are indicated in red, yellow, and blue, respectively. The amino acid residues mutated by 7F4 selective pressure are highlighted in the magenta circles. (B) Positions of the amino acid mutations on a DENV prM-E protein heterodimer. The ribbon diagram is based on PDB accession No. 3C5X. prM protein is indicated in green.

**Table 2.** Summary of the escape mutants.

|  | Cell | prM-13 | E-58 | E-67 | E-69 | E-118 | NT<sub>50</sub><br>(µg/mL) | Number<br>of glycans |
| --- | --- | --- | --- | --- | --- | --- | --- | --- |
| Mochizuki WT | - | M | E | N | I | K | 0.02 | 1 |
| Group 1 | Vero | I | E | N | I | E | >29 | 1 |
| Group 2 | Vero | I | E | D | I | K | >29 | 1 |
| Group 3 | Vero | I | K | N | T | K | >29 | 2 |
| Group 4 | C6/36 | I | E | N | T | K | >29 | 2 |

The N-linked glycosylation motif was N<sub>67</sub>-X<sub>68</sub>-T/S<sub>69</sub>.

### Determination of 7F4 epitope by site-directed mutagenesis

The mutations at M13I in the prM and E58K in the E were located relatively far from the N-67 glycosylation site, compared with the other mutations (N67D, I69T, and K118E in the E) (Fig. 2). Furthermore, the 13^th^ amino acid in the prM (prM-13) was not included in the mature infectious virion because the pr protein was cleaved through the maturation step. Moreover, E58K was not introduced in the escape mutant derived from the C6/36 cells. Thus, we hypothesized that prM-13 and E-58 were not involved in the 7F4 epitope, but with concomitant mutations to the other mutations. Therefore, we examined whether or not those mutations affect reactivity of 7F4 by constructing mutant virus-like particles (VLPs).

The single mutation at M13I in the prM and E58K in the E did not affect 7F4 reactivity, although other mutations affected it (Fig. 3A). Even double mutations at those residues did not affect it, indicating that prM-13 and E-58 were not a part of the 7F4 epitope, but considered as concomitant mutations. Conversely, E-67, E-69, and E-118 were confirmed to be 7F4 epitopes.

**Figure 3.**
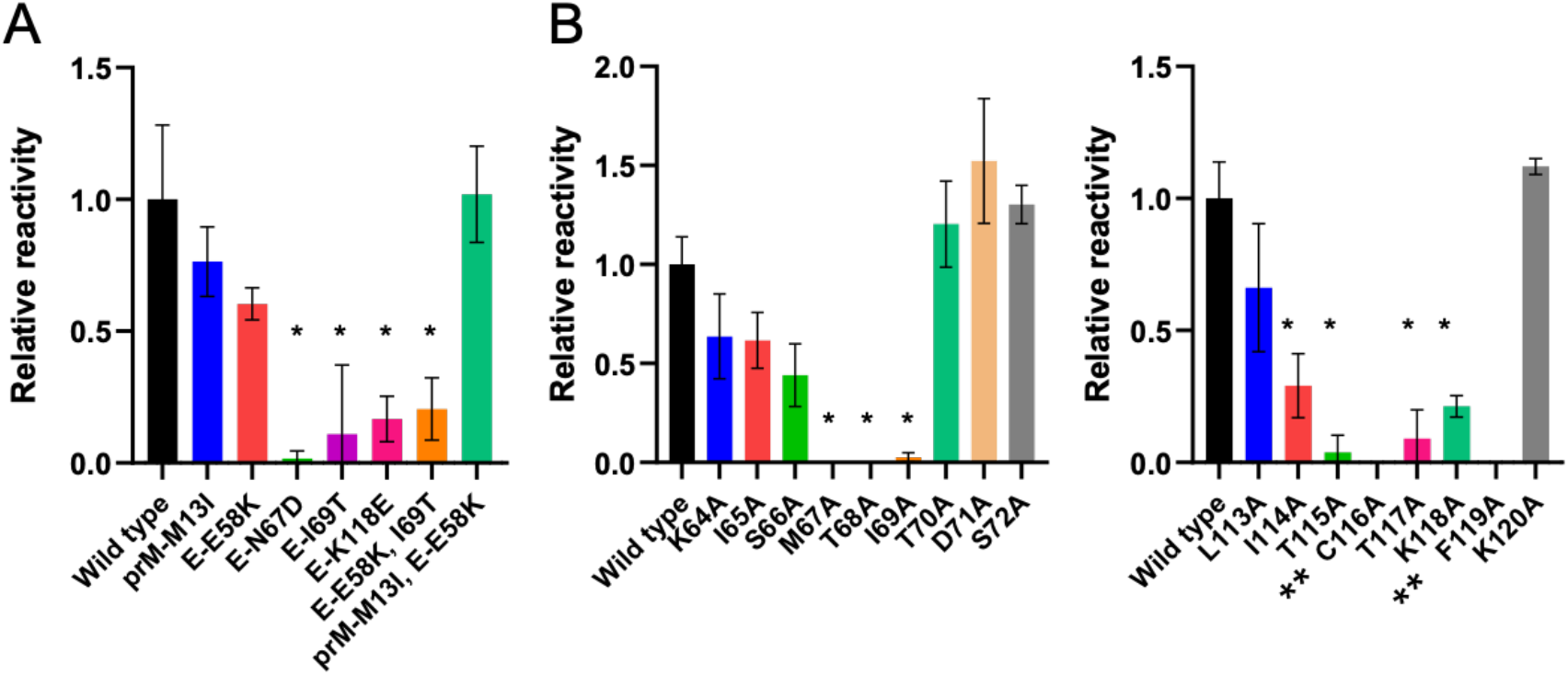
Epitope mapping by site-directed mutagenesis. (A) Reactivity of 7F4 to the mutant VLPs. VLPs were produced from the 293T cells transduced with prM-E expression plasmids with or without mutations. Reactivity was expressed as the ratio of 7F4 to 4G2. Reactivity to wild type VLP was set to 1.0. The averages and SDs of the two independent experiments are shown. Detergent was not added in this ELISA system to maintain viral particle form. A one-way ANOVA was performed to analyze statistical significance comparing wild type and mutants. * P < 0.05. (B) Alanine scanning. Averages and SDs of the two independent experiments are shown. A one-way ANOVA was performed to analyze the statistical significance comparing wild type and mutants. * P < 0.05. **C116A and F119A mutants were not detected by 4G2, indicating inefficient VLP release from the transduced cells.

To narrow down the detailed epitope, alanine substitutions were introduced into the positions near the deduced epitope; E-64 to E-72, and E-113 to E-120. Mutant VLPs with alanine substitution at E-67 to E-69 and E-114 to E-118 lost their reactivity to 7F4 (Fig. 3B), thus indicating that those residues were involved in the 7F4 epitope.

### Determination of 7F4 epitope by comparing the Mochizuki and Hawaii strains

The Mochizuki strain was neutralized by 7F4, whereas the Hawaii strain was not neutralized. There are 10 differences between the amino acids of the Mochizuki and Hawaii strains in the ectodomain of E (Fig. 4A). Out of these, amino acid differences at E-69 and E-117 may render Hawaii strain nonreactive to 7F4 because these residues were included in the deduced 7F4 epitope (Fig. 4B). Thus, we introduced T69I and A117T mutations into the Hawaii strain in order to conform the Hawaii strain to Mochizuki strain in terms of the 7F4 epitope. However, mutant Hawaii VLPs were still nonreactive to 7F4 (Fig. 4C). Therefore, we next added Mochizuki-like mutations into three more residues (Q228K, E234Q and V251A in the E) in the Hawaii strain because these residues were structurally near the N-67 glycan (Fig. 4B). As a result, four simultaneous mutations (T69I, A117T, E234Q, and V251A) made mutant Hawaii VLP reactive to 7F4, at a level similar to that of the Mochizuki strain (Fig. 4C).

**Figure 4.**
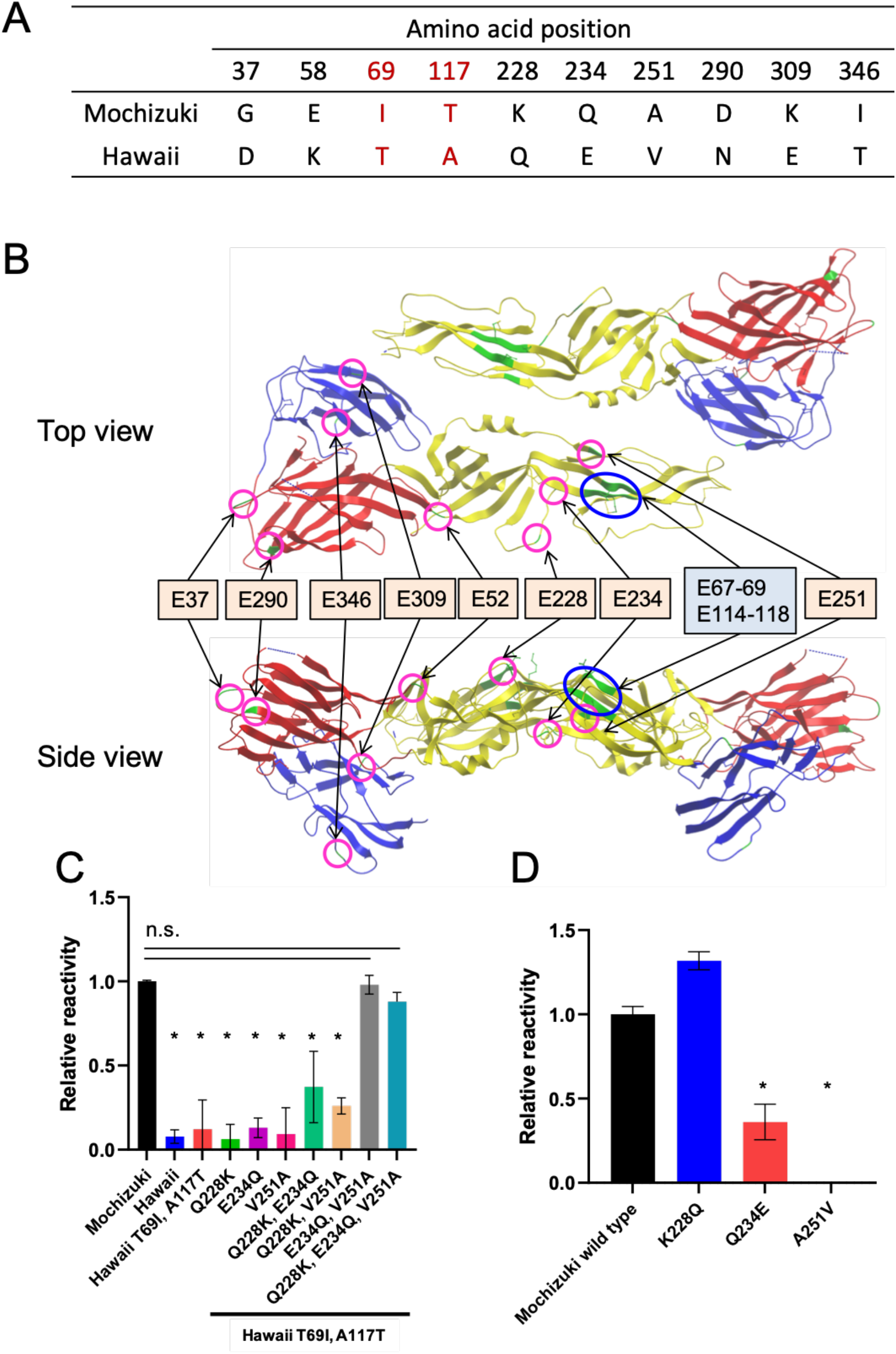
Comparison of Mochizuki and Hawaii strains. (A) Amino acid differences between the Mochizuki and Hawaii strains in the E protein. (B) Positions of the amino acid differences in the E-dimer. The ribbon diagram is based on PDB accession No. 4GSX. EDI, EDII, and EDIII are indicated in red, yellow, and blue, respectively. Amino acid differences between Mochizuki and Hawaii are highlighted in the magenta circles. 7F4 epitope is highlighted in the blue circle. (C) Reactivity to the mutant Hawaii VLPs. Reactivity was expressed as the ratio of 7F4 to 4G2. The reactivity to wild type VLP was set to 1.0. Averages and SDs of the two independent experiments are shown. A one-way ANOVA was performed to analyze the statistical significance comparing wild type Mochizuki and mutants. * P < 0.05. n.s. not significant (P > 0.05). (D) Reactivity to mutant Mochizuki VLPs.

Finally, we introduced three Hawaii-like mutations into the Mochizuki strain (K228Q, Q234E, and A251V). Q234E and A251V mutants lost their reactivity to 7F4 (Fig. 4D), thus validating that those two residues (E-234 and E-251) were also involved in the 7F4 epitope.

### Growth curve of the escape mutants

The Mochizuki strain was very unique in E at E-58 and I at E-69, compared to naturally circulating DENV-1 (Table S1). Interestingly, the selective pressure of 7F4 mutated these unique amino acid residues to those of naturally circulating dominant DENV-1 strains (i.e. E-E58K and E-I69T observed in the group 3 escape mutant). We hypothesized that the group 3 mutant had superior viral fitness compared to the other mutants. Thus, we further analyze the impact of the mutations on viral replication using growth curve assays (Fig. 5). The mutant with K118E (group 1) showed the same high level of viral growth as WT Mochizuki, except for day 5 in Vero cells. The mutant with N67D (group 2) showed 1,000-fold lower virus replication than the others in Vero cells, while higher viral replication was seen in C6/36 cells. The mutant with only I69T (group 4) showed lower replication than that of WT Mochizuki in both Vero and C6/36 cells. However, the mutant with E58K/I69T (group 3) showed the same high level of viral growth as the WT Mochizuki strain, indicating that E58K rescued the growth of glycan-acquired mutant.

**Figure 5.**
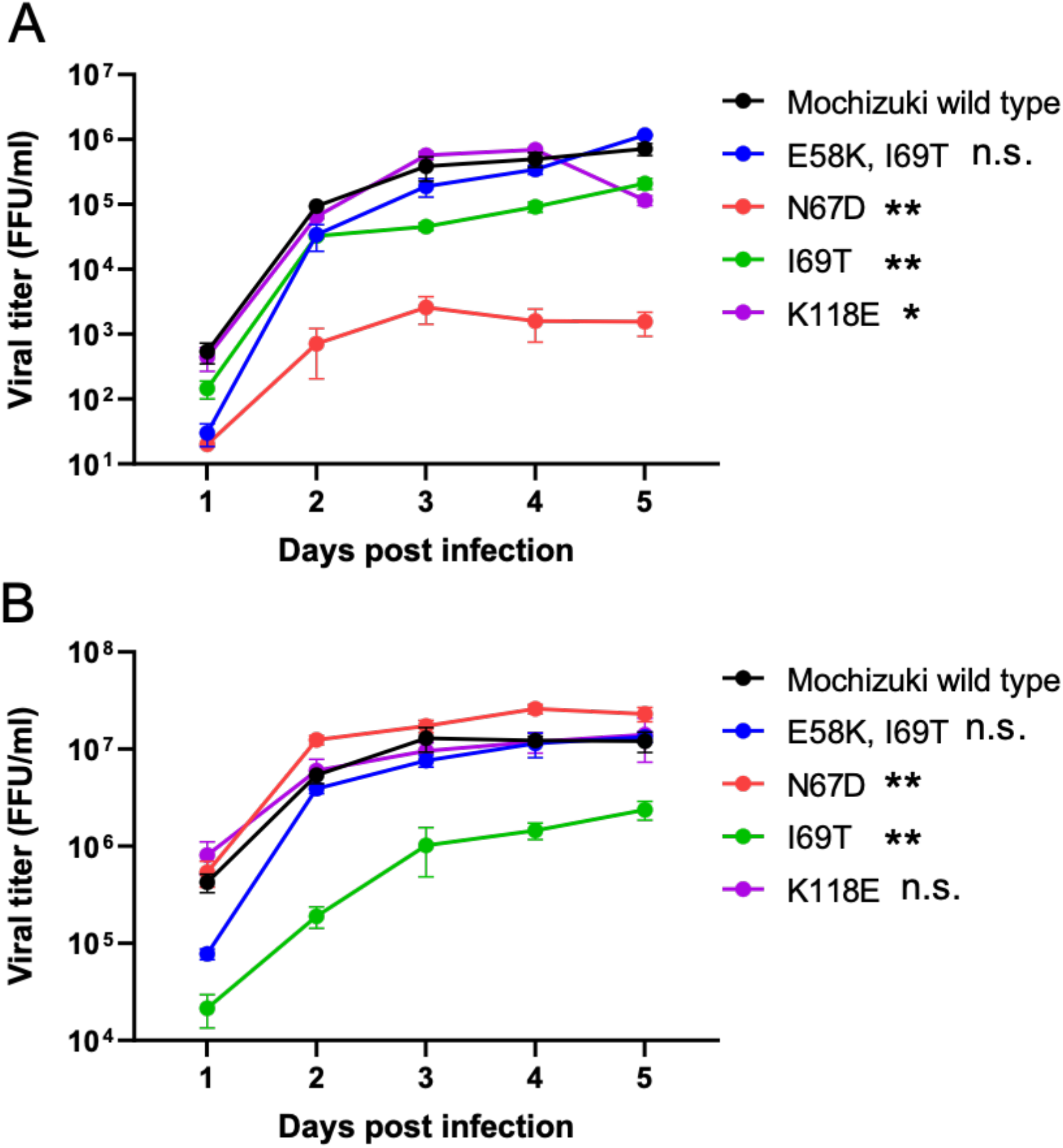
Viral growth curve assay using the escape mutants. (A) Growth curve using Vero cells. MOI was set to 0.01. Averages and SDs of the two independent experiments are shown. A two-way ANOVA was performed to analyze the statistical significance comparing wild type and mutants. * P < 0.05. ** P < 0.01. n.s. not significant (P > 0.05). (B) Growth curve using C6/36 cells. MOI was set to 0.01.

### Effect of deglycosylation on the reactivity to 7F4

Acquisition of N-67 glycan in the 7F4 escape mutants suggested another function of N-67 glycan as a shield against neutralizing antibodies, as reported in HIV-1 or influenza virus (Crispin et al., 2018; Kobayashi and Suzuki, 2012). Therefore, we hypothesized that deglycosylation may recover 7F4 reactivity to the escape mutants with I69T. 7F4 reactivity in the original and deglycosylated escape mutants was compared using ELISA. However, there was no significant difference between the original and deglyscosylated viruses, suggesting that N-67 glycan was not related to antibody binding (Fig. 6). Amino acid sequence (N_67_-I_68_-I_69_) was important rather than glycosylation.

**Figure 6.**
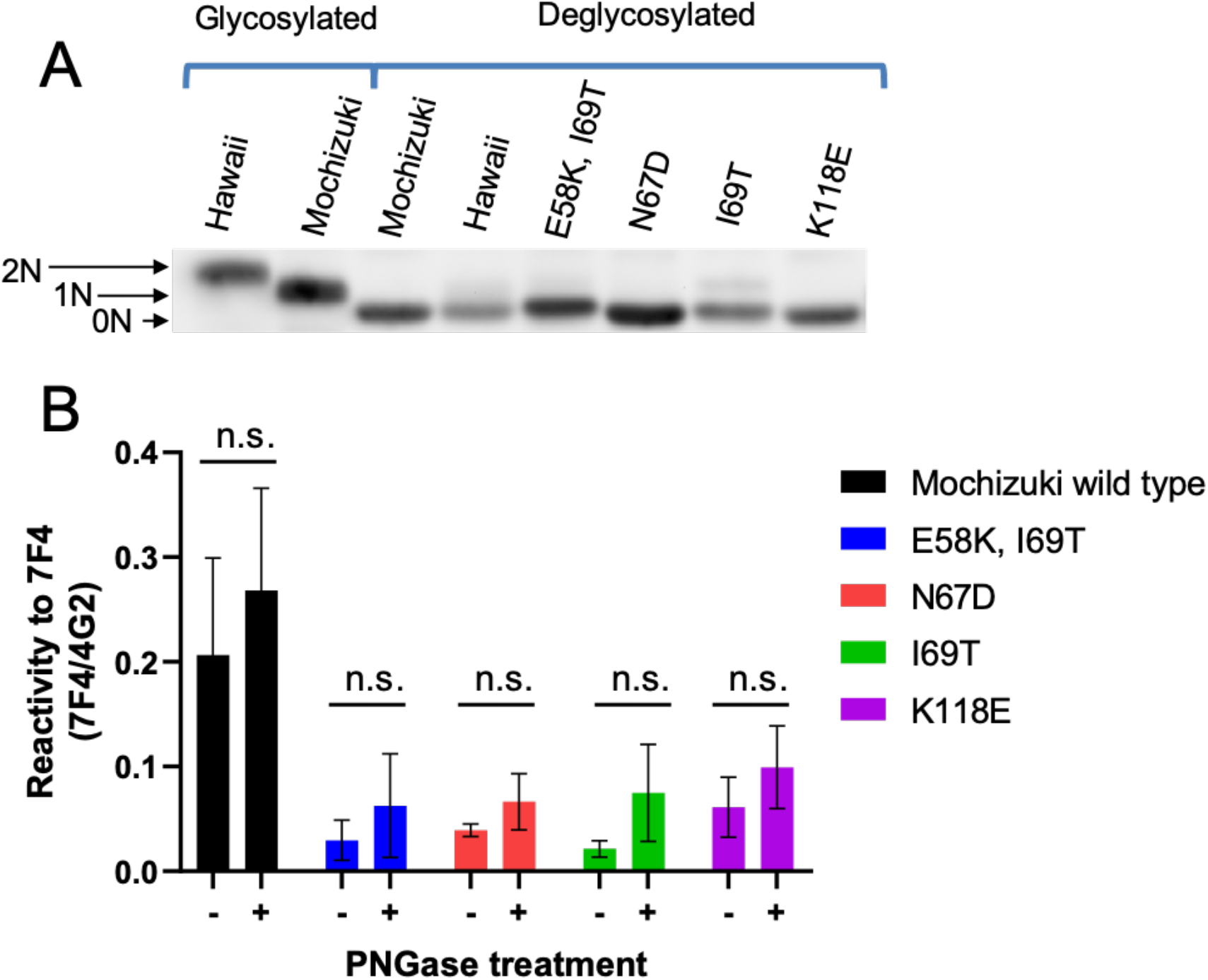
Comparison of 7F4 reactivity of glycosylated and deglycosylated viruses. (A) Confirmation of deglycosylation by Western blot. The E protein was denatured under nonreducing conditions and separated in SDS-polyacrylamide gel. The glycan number is shown with arrows. (B) Reactivity of 7F4 to the glycosylated and deglycosylated viruses determined with ELISA. Reactivity to 7F4 was expressed as the ratio of the OD of 7F4 and 4G2. Averages and SDs of the two independent experiments are shown. Multiple t-tests were conducted to analyze statistical significance comparing glycosylated and deglycosylated viruses. n.s. not significant (P > 0.05).

### Animal experiments

Considering that all currently circulating DENVs possess the N-67 glycan, this epitope is no longer useless as a vaccine target. Nevertheless, antibodies competing for 7F4 were detected in the serum of humans infected with DENV (Yamanaka et al., 2013). Therefore, we analyzed whether or not ablation of the 7F4 epitope disrupts immunogenicity to elicit 7F4-like antibodies. prM-E expression plasmids encoding WT Mochizuki, N67D mutant, and I69T mutant were immunized to BALB/c mice. NT_50_ of the mouse sera immunized with WT, N67D, and I69T were 1:40, 1:40, and 1:20, respectively. A competition assay using biotinylated 7F4 revealed that the proportion of 7F4-like antibody was decreased in N67D and I69T immunized mice, as expected (Fig. 7). However, induction of 7F4-like antibodies appeared to be not completely suppressed by those mutations, although reactivity was completely disrupted. This suggested that there are epitopes competing for 7F4 epitope on the E protein.

**Figure 7.**
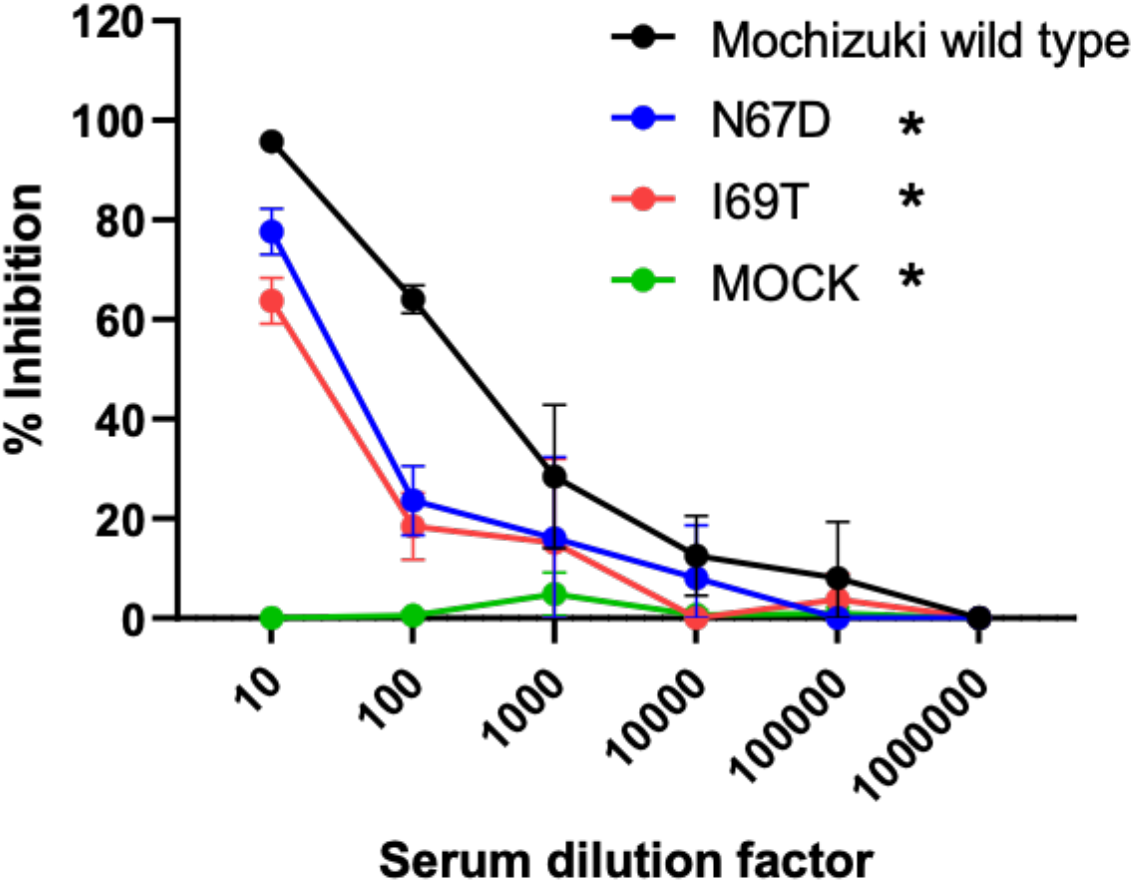
Competition assay using biotinylated 7F4 and immunized mouse sera. Averages and SDs of the two independent experiments are shown. A two-way ANOVA was performed to analyze statistical significance comparing wild type and mutants. * P < 0.05.

## 3. Discussion

Understanding the neutralizing mechanism of DENV infection will be useful in the development of an effective vaccine against dengue, which has not yet been available. The epitope of highly potent neutralizing antibody called 7F4 was further analyzed in this investigation, by means of escape mutant and site-directed mutagenesis. In summary, amino acid residues E-67 to E-69, E-114 to E-118, E-234, and E-251 were presumably involved in the 7F4 epitope (Figs. 3, 4). The epitope was located at the region close to the N-67 glycan on the DII. The glycan did not function as an antibody shield, as shown in Fig. 6. This epitope is distinct from the previously reported neutralizing epitopes (Teoh et al., 2012; de Alwis et al, 2012; Dejnirattisai et al, 2015). Several potent E-dimer epitope antibodies partly target the epitope near the N-67 glycan, suggesting the potency of the epitope around N-67 glycan (Roubinski et al., 2015). This investigation demonstrated the usefulness of the epitope on the central region of DII for the development of a dengue vaccine. DII is still an important immunogen even though DIII subunit vaccine showed promising results (Block et al., 2010; Park et al., 2020).

7F4 recognized E-monomer epitope because 7F4 can react to detergent-inactivated E protein determined by ELISA. Notably, 7F4 inhibited both pre- and postadsorption steps despite its E-monomer recognition (Fig. 1). 7F4 binds to the lateral part of E protein such as E-234 and E-251, which are hidden by the adjacent E protein when it forms E-dimer (Figs. 2, 4). 7F4 might bind to this cryptic epitope during viral breathing, resulting in the block of viral conformational change. Alternatively, the antibody efficiently binds to the E-trimer during membrane fusion. Further analysis on the neutralization mechanism is needed.

7F4 selective pressure changed the unique amino acid residues in the Mochizuki strain to those of currently circulating DENV-1 (Tables 2, S1). Viral growth curve assays showed that E58K/I69T and K118E mutants showed viral replication rates comparable to that of WT Mochizuki, while N67D and I69T mutants showed less viral propagation in Vero and/or C6/36 cells (Fig. 5). Acquisition of N-67 glycan is reportedly beneficial for viral attachment to monocyte/macrophage cells and virion release (Mondote et al., 2007; Lee et al., 2010). These results demonstrated why E58K/I69T becomes dominant in nature. There is a limitation that viral growth curve assays were conducted directly using the escape mutants because the infectious clone of the Mochizuki strain was not available. Thus, prM-13 mutation introduced in all the escape mutants might have affected the viral growth. Nevertheless, the results still demonstrated the best fitness of E58K/I69T mutant among the escape mutants. This antibody selection pressure might facilitate viral evolution.

As shown above, 7F4 was absolutely specific to the Mochizuki strain and cannot react to most currently circulating DENVs. However, 7F4-like antibodies were induced in individuals infected with currently circulating DENVs (Yamanaka et al., 2013). This study demonstrated the existence of epitopes competing for 7F4 epitope. The antibodies competing for 7F4 epitope played an important role in neutralization in humans since their amounts were correlated with the neutralizing titer (Yamanaka et al., 2013). This suggested the importance of the induction of antibodies recognizing epitopes near the N-67 glycan for future dengue vaccine development.

In conclusion, we identified a novel potent neutralizing epitope near the N-67 glycan on the DII of the E protein. This information would be helpful for the development of therapeutic antibody or vaccine candidates against dengue.

## Materials and Methods

### Cells, viruses, and antibodies

Vero cells were cultured in Eagle’s minimum essential medium (MEM) supplemented with 10% fetal bovine serum (FBS) at 37°C in 5% CO_2_ and 95% air (Konishi and Fujii, 2002). The C6/36 cells were cultured in MEM supplemented with 10% FBS and nonessential amino acids at 28°C in 5% CO_2_ and 95% air (Konishi and Fujii, 2002). The HEK-293T cells were maintained in Dulbecco’s modified Eagle medium supplemented with 10% FBS.

The Mochizuki and Hawaii (Genbank accession number KM204119) strains, and an Indonesian clinical isolate (A4 strain: Genbank accession number AB600922) of DENV-1, were used in this study (Konishi et al., 2006; Sjatha et al., 2012). Viruses harvested from culture fluids of infected C6/36 cells were used for subsequent experiments. 7F4, a mouse MAb derived from Mochizuki strain immunized mouse, was used (Yamanaka et al., 2013). Furthermore, 4G2 (E specific, cross-reactive to flavivirus group) was used for the detection of virus-infected cells (American Type Culture Collection, Manassas, VA).

### Titration of viral infectivity and neutralization test

Infective titers were determined on Vero cells by counting the infectious foci after immunostaining (see below) and expressed as focus-forming units (FFU).

A neutralizing test was performed as described previously (Konishi et al., 2006). Briefly, Vero cells were seeded in a 96-well plate (2 × 10^4^ cell/well). The next day, 100 FFUs of viruses and the serially diluted antibody were incubated for 1 hour and then inoculated onto the cells. The cells were then incubated overnight at 37°C, followed by immunostaining and focus counting. The neutralizing antibody titer was expressed as the maximum IgG concentration or serum dilution yielding a 50% reduction in focus number (NT_50_).

### Time of addition assay

Time of addition assays were performed as described previously (Hung et al., 1999). For the preadsorption assay, the virus was preincubated with serially diluted antibodies at 4°C for 1 hour and then inoculated onto Vero cells seeded, as described in the neutralization test. Then, the unadsorbed viruses and excess antibodies were washed out with PBS. The cells were then incubated overnight at 37°C, followed by immunostaining and focus counting.

For the postadsorption assay, the virus was added directly to cells at 4°C for 1 hour. The unadsorbed viruses were removed by washing the cells, and the bound viruses were incubated with serially diluted antibodies at 4°C for one additional hour. The cells were then incubated at 37°C overnight, followed by immunostaining and focus counting. The results were expressed in the same manner as in the neutralization assay.

### Immunostaining

Immunostaining was performed as previously described (Konishi and Fujii, 2002). Briefly, infected cells were fixed with acetone-methanol (1:1). These cells were incubated serially with 4G2, biotinylated anti-mouse IgG, VECTASTAIN Elite ABC kit, and VIP peroxidase substrate kit (Vector Laboratories, Burlingame, CA).

### Generation of the neutralization-escape mutants of the Mochizuki strain to 7F4

Mochizuki strain was inoculated onto Vero or C6/36 cells and incubated for 1 hour to allow for viral uptake. The cells were washed three times with PBS and then cultured with MEM containing 7F4 at various concentrations (29 μg/mL, 23.2 μg/mL, 11.6 μg/mL, 5.8 μg/mL, 3.6 μg/mL, and 2.9 μg/mL). Culture supernatants were transferred to the new cells after 7 days of incubation. These processes were repeated five times. Passage control strain was generated as well.

### RNA extraction, RT-PCR, and sequence analysis

Viral RNA was extracted from the culture supernatant of the escape mutant infected cell using TRIzol Reagent (Invitrogen, Carlsbad, CA). RT-PCR was conducted using the Thermoscript RT-PCR system (Invitrogen, Carlsbad, CA) and Ex Taq (Takara, Shiga, Japan) to amplify the prM-E coding regions. A gene-specific sense primer designed in the C coding region [5’-GGGAGAGTTATGTGAGG-3’; corresponding to nucleotides 559-575 of the Mochizuki strain (Genbank accession No. AB074760)] and an antisense primer designed in the nonstructural 1 (NS1) coding region (5’-CCCAGCTTTTCCACGAG-3’: nucleotides 2,774-2,758) were used.

### Plasmid construction and VLP preparation

The pcDNA3 plasmids encoding the prM-E of the DENV-1 of the Mochizuki strain (designated as pcD1ME) or Hawaii strain (pcD1HME) were constructed following a previous report (Konishi et al., 2006). Then, mutations were introduced into the plasmids using a site-directed mutagenesis kit following the manufacturer’s protocol (Toyobo, Osaka, Japan).

The plasmids were transfected into the 293T cells using Fugene HD transfection reagent (Promega, Madison, WI). The culture media containing VLP were harvested 3 days post transfection. The harvested VLP was subjected to determination of reactivity to 7F4 using ELISA (see below).

### Determination of the reactivity of 7F4 to mutant VLPs

Rabbit serum immunized with DENV-1 Mochizuki strain (in-house product) was coated onto a Nunc Maxisorp ELISA plate (Thermo Fisher Scientific, Waltham, MA). Then, the antigens (Mochizuki, Hawaii, or mutant VLPs), first antibody (4G2 or 7F4), alkaline phosphatase-conjugated anti-mouse IgG (Jackson ImmunoResearch Laboratories, West Grove, PA), and p-nitrophenyl phosphate (Nacalaitesque, Kyoto, Japan) were serially incubated, followed by measurement of the absorbance at 415 nm. The ELISA diluent was composed of PBS containing 1% bovine serum albumin and 0.05% Tween 20, unless otherwise mentioned. The reactivity to 7F4 was expressed as relative value of 7F4 to 4G2 (OD of 7F4/OD of 4G2) in order to standardize the amount of VLP. The introduced mutations did not affect the epitope of 4G2 [fusion loop (98^th^-110^th^ amino acids) of the E protein]. The values obtained from WT VLP (without mutation) were converted to 1.0, and the values of mutant VLPs were converted in the same way.

### Growth curve of the escape mutants

Vero or C6/36 cells were seeded in a 6-well plate (5 × 10^5^ cell/well). Mochizuki strain or the escape mutants were infected to the cells at multiplicity of infection (MOI) of 0.01 the next day. The cells were washed with PBS three times after one hour of incubation. Two mL of medium was added to the well, and the plate was incubated at 37°C. The culture supernatants were collected every 24 hours post infection. The viral infective titers of the supernatants were determined on Vero cells.

### Retrieval of DENV-1 E sequences

The E sequences showing similarity to the Mochizuki strain were retrieved by BLAST. A total of 4,916 sequences were retrieved and aligned using Genetyx ver. 10 (Genetyx, Tokyo, Japan). The amino acid differences at E-58, E-67, E-69, and E-118 were examined manually. The data set is available upon request.

### Deglycosylation of E protein

Deglycosylation of the DENV was performed using PNGase F (New England Biolabs, Ipswich, MA) and Endo H (New England Biolabs, Ipswich, MA), following the manufacturer’s protocol. Deglysosylated samples were subjected to Western blot to confirm deglycosylation (see below). Furthermore, deglycosylated E protein was examined in its reactivity to 7F4 by ELISA.

### Western blot

Western blot was performed as previously described (Konishi et al., 1992). Briefly, the samples were denatured by heating them at 95°C for 5 minutes under reducing or nonreducing conditions, and run on 9% SDS-polyacrylamide gel electrophoresis (SDS-PAGE), followed by transfer to polyvinylidene difluoride membranes (Millipore Corporation, Bedford, MA). E protein was detected through serial incubation of detector antibody (4G2 or 7F4), HRP-conjugated anti-mouse IgG (Promega, Madison, WI), and ImmuonStar Zeta substrate (FUJIFILM Wako Pure Chemical Corporation, Osaka Japan).

### Animal experiments

Three kinds of plasmids (pcD1ME, pcD1ME with N67D, and pcD1ME with I69T) were immunized to 4-week-old male C3H/He mice (SLC Japan, Shizuoka, Japan). Groups of six mice were immunized with 50 μg of plasmid DNAs via intracutaneous route three times at an interval of 2 weeks. The blood was collected from the immunized mice 1 week after the third immunization. The pooled sera were subjected to a neutralizing test as well as a competition assay.

### Competition assay

The competition assay was performed according to a previously described method (Yamanaka et al., 2013). Briefly, ELISA plates coated with rabbit hyperimmune sera against DENV-1 were serially incubated with Mochizuki strain, serially diluted immunized sera, biotinylated 7F4, avidin-alkaline phosphatase conjugate, and p-nitrophenyl phosphate. The biotinylated antibody was prepared using a purified IgG fraction of MAb and EZ-Link Sulfo-NHS-LCBiotin [sulfosuccinimidyl6(biotinamido)hexanoate] (Pierce, Rockford, IL) according to the manufacturer’s instructions. Percentages of competition were calculated by comparing the sample OD with that obtained without incubation with the immunized sera.

### Statistics

All error bars indicated standard deviations. All the calculations were performed using GraphPad Prism 8 (GraphPad Software Inc.).

## Acknowledgements

This work was supported by a program from the Japan Initiative for Global Research Network on Infectious Diseases (J-GRID) through the Ministry of Education, Culture, Sport, Science and Technology in Japan, and the Japan Agency for Medical Research and Development (AMED); KAKENHI Grant-in-Aid for Young Scientists (17K16224; 20K20200). The BIKEN Endowed Department of Dengue Vaccine Development was established by an endowment from the Research Foundation for Microbial Diseases of Osaka University, Osaka, Japan, to the Research Institute for Microbial Diseases, Osaka University, Osaka, Japan. The funders had no role in study design, data collection and analysis, decision to publish, or preparation of the manuscript.

## Author contributions

All authors conceived the study. T.K. performed the experiments and took the lead in writing the manuscript. A.Y., E.K. and M.K. provided feedback and helped shape the research and manuscript.

