## Supplementary information for "A potent neutralizing mouse monoclonal antibody specific to dengue virus type 1 Mochizuki strain recognized a novel epitope around the N-67 glycan on the E protein: a possible explanation of dengue virus evolution regarding the acquisition of N-67 glycan"

**Table S1. Comparison of amino acid sequences on the E region among Mochizuki, escape mutant, and naturally circulating DENV-1**

| | Envelope region | | | |
|---|---|---|---|---|
| | 58 | 67 | 69 | 118 |
| Mochizuki | E | N | I | K |
| Escape mutant | K | D | T | E |
| | K<br>4912<br>(99.92%) | N<br>4901<br>(99.70%) | T<br>4913<br>(99.94%) | K<br>4911<br>(99.90%) |
| | R<br>2<br>(0.04%) | D<br>14<br>(0.28%) | I<br>3<br>(0.06%) | E<br>2<br>(0.04%) |
| Other DENV-1 strains (4916 strains) | I<br>1<br>(0.02%) | K<br>1<br>(0.02%) | | R<br>1<br>(0.02%) |
| | E<br>1<br>(0.02%) | | | M<br>1<br>(0.02%) |
| | | | | N<br>1<br>(0.02%) |

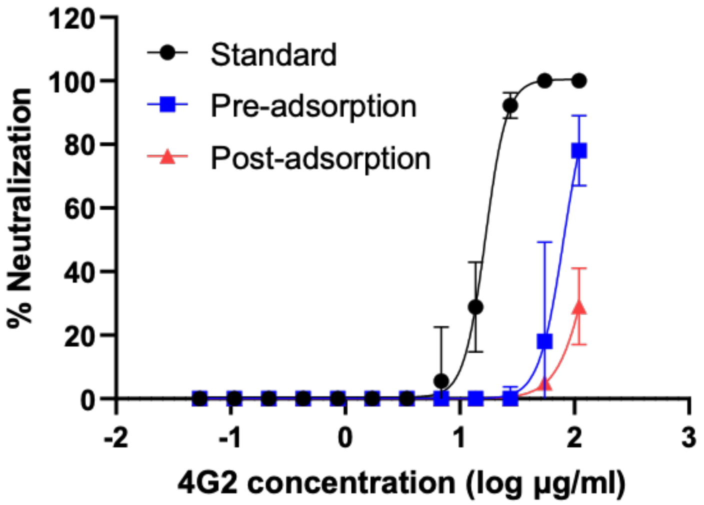


**Figure S1. Time of addition assay of 4G2**

Averages and SDs of the two independent experiments are shown.

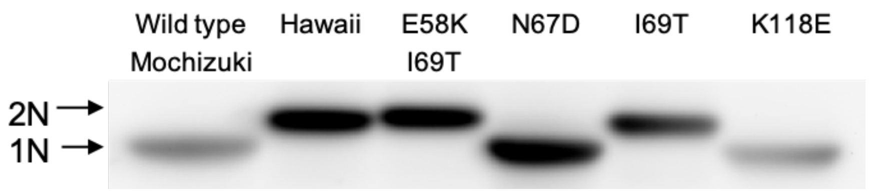


**Figure S2. Confirmation of the number of glycan in the E protein**

The E protein was denatured under nonreducing conditions and separated on an SDS-polyacrylamide gel. The upper size band indicates two glycans, while the lower band indicates one glycan.
